# Action inhibition revisited: a new method identifies compromised reactive but intact proactive control in Tourette disorder

**DOI:** 10.1101/2021.06.15.448397

**Authors:** Indrajeet Indrajeet, Cyril Atkinson-Clement, Yulia Worbe, Pierre Pouget, Supriya Ray

## Abstract

Tourette disorder (TD) is characterized by tics, which are sudden repetitive involuntary movements or vocalizations. Deficits in inhibitory control in TD patients remain inconclusive from the traditional method of estimating the ability to stop an impending action, which requires careful interpretation of a parameter derived from race model. One possible explanation for these inconsistencies is that race model’s assumptions are often violated. Here, we used a pair of metrics derived from a recent alternative model to address why stopping performance in TD patients is unaffected by impairments in neural circuitry. These new metrics distinguish between proactive and reactive inhibitory control and estimate them separately. When these metrics were contrasted with healthy controls (HC), we identified robust deficits in reactive control in TD patients, but not in proactive control. The patient population exhibited difficulty in slowing down the speed of movement planning, which they compensated by their intact ability of procrastination.

**TEASER:** Tourette disorder patients inhibit action by means of strategic postponement to compensate impaired slowness in preparation.

## INTRODUCTION

The ability to inhibit unwanted movements is one of the prerequisites for flexible goal-directed behavior. In the laboratory, inhibitory control is often assessed using standard tasks like Go/No-Go and countermanding (or stop-signal) tasks. In Go/No-Go task, the commission error rate (i.e., the frequency of errantly eliciting a response when participants are not supposed to do so) is used as a metric of proactive inhibitory control. In the countermanding task, race model (*1*) describes performance as an outcome of a competition between a preparatory GO and an inhibitory STOP process, each rising independently at a stochastic rate to reach a common fixed threshold. Accordingly, the model estimates the average time taken by STOP process to reach the threshold, called stop-signal response time (SSRT), which is widely used as a metric of inhibitory control. The shorter the SSRT, the better the inhibitory control.

The average SSRT in patients population has been contrasted with healthy control (HC) to identify the propensity of impairment in inhibitory control in the populations with motor and cognitive deficits like Parkinson’s disease (*2, 3*), schizophrenia (*4*–*6*), obsessive-compulsive disorder (*7, 8*), attention-deficit hyperactivity disorder (*9, 10*), and Tourette disorder (TD) (*11, 12*). Tourette disorder (TD) represents a relevant model of inhibition impairment due to its major clinical sign: tics. Usually, they corresponded to sudden, repetitive, non-rhythmic, involuntary or semi-voluntary movements and/ or vocalization (*13, 14*) and are frequently considered as *“fragments of motor behavior that escape voluntary motor control”*, related to a deficiency in inhibitory control of actions (*11, 15, 16*). However, studies did not find a consistent significant difference between TD and HC either in the commission error rates (e.g., *8, 17, 18*; *but also see 19*) or in SSRT (e.g., *11, 20*). One possible explanation for the inconsistent findings of difference in SSRT is that SSRT might not be an appropriate measure of inhibitory control when the context independence assumption (i.e., STOP process does not influence GO process) is violated, especially at short SSDs (*21, 22*). The violation is pervasive and has a severe effect on the estimation of SSRT. For example, if trials with short intervals between the go and stop signal are removed from SSRT estimation, the difference reported in published studies between HC and disorder groups mostly disappears (*23*–*25*).

To assess the contributions of both proactive and reactive control in movement inhibition without independence assumption, we derived a pair of metrics on the foundation of the cancellable rise-to-threshold (CRTT) framework of countermanding, which we referred to as ‘proactive delay’ and ‘log-attenuation rate’, respectively (*22, 26, 27*). The former estimates the time consumed in procrastinating the response in expectation of the stop-signal to increase the odds of successful inhibition, whereas the latter estimates the dynamics of decelerated preparatory build-up activity triggered by the onset of the stop-signal. Since CRTT metrics are independent of the race model’s assumptions and sensitive to deficits in proactive and reactive inhibitory control independently, we revisited the deficits in inhibitory control in TD patients (*12*).

## RESULTS

A total of 63 TD and 34 HC performed an Emotional Stop Signal Task (ESST), which required them to refrain from pressing a key in response to an infrequent stop-signal (*12*) (see Figure 1 & materials and methods)

**Figure. 1.**
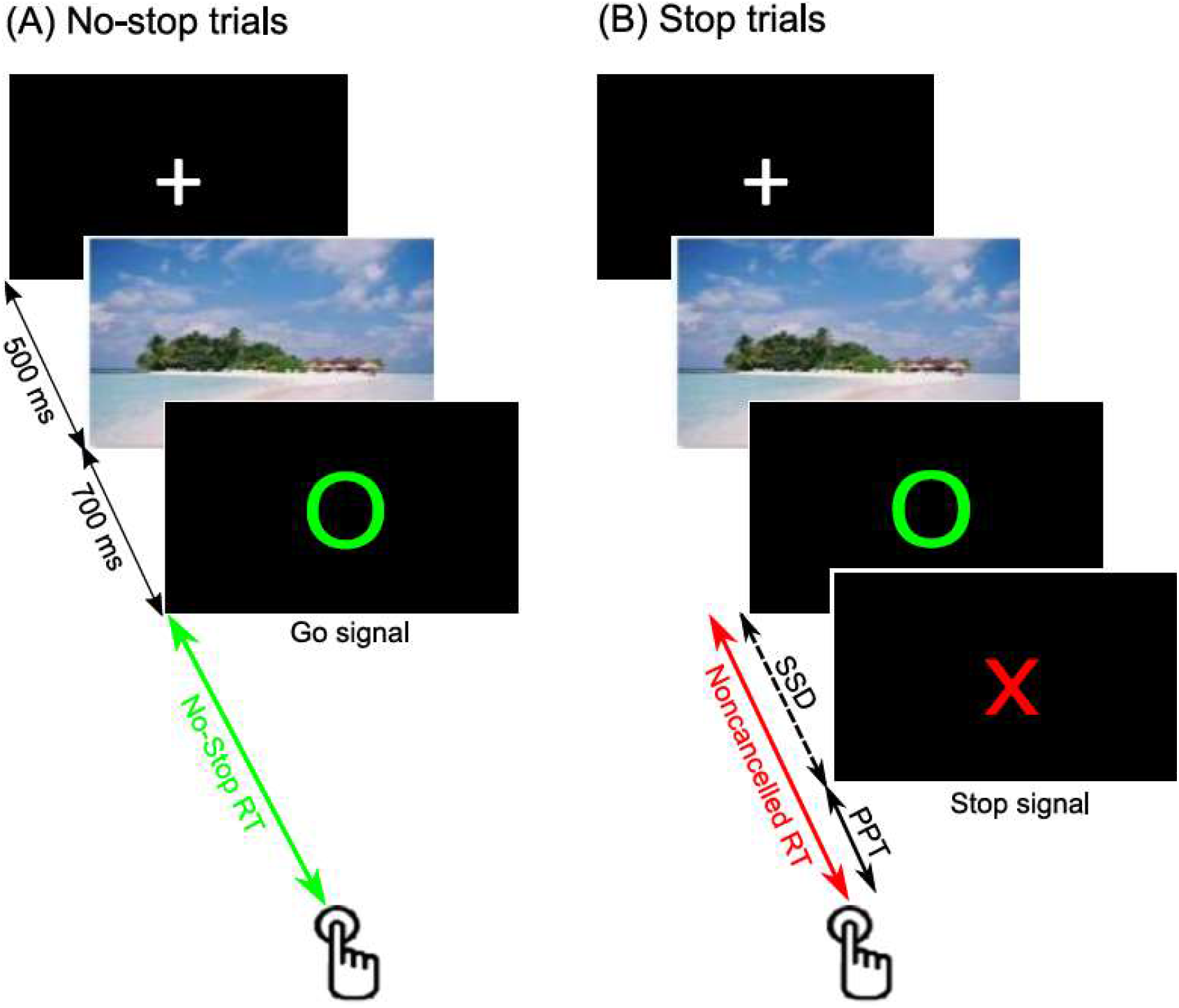
Schematic of Emotional Stop Signal Task (ESST). Each trial began with the display of a fixation cross (+) in white color at the center of the display. After 500 ms, an image of a scene with emotional content appeared at the center for 700 ms. Subsequently, a blank display in black appeared for a variable 1-2 s interval. It was followed by a green unfilled circle at the center. **(A)** In the majority (66)%of trials (no-stop trial), participants were instructed to press the ENTER key on the keyboard “*as soon as psosible*”. **(B)** In the rest of the trials (stop trials), after a variable delay from the circle, the letter X in capital in red color appeared at the center, instructing participants to refrain from pressing the ENTER key. No-stop and stop trials were pseudorandomly presented.

### Participant’s demographics

The data collected from 14 participants (10 TD patients and 4 HC) were incomplete, hence could not be used for analyses. A total of 53 TD (Sex: 11 females, 42 males, mean ± SD age: 30.21 ± 10.5 years, education: 14.25 ± 2.56 years) and 30 HC (Sex: 09 females, 21 males, mean ± SD age: 31.63 ± 10.44 years, education: 14.57 ± 2.93 years) was available for this study. In this sample, there was no significant difference in age [*t* (81) = 0.596, *p* (two-tailed) = 0.553, *d* = 0.136, BF_01_ = 3.624]; education level [*t* (81) = 0.522, *p* (two-tailed) = 0.603, *d* = 0.119, BF_01_ = 3.756]; or gender [*X*^2^(1, *N* = 83) = 0.895, *p* = .344, Cramer’s *V* = 0.012] between the TD and HC groups. Out of 53 TD patients, 19 were taking antipsychotic medication, 23 had ADHD, and 11 had OCD. The mean (SD) tic severity was 16.321(7.076) as measured by YGTSS/50 scale. After removing the outliers (see supplementary materials for details), the final sample consists of 28 HC and 49 TD participants.

### Behavioral performances

No significant differences were found between HC and TD on all behavioral performances (i.e., Go accuracy, error rate, SSD, no-stop RT, and stop error RT; see Table 1). First, we binned the SSD from 0 to 600 ms in the size of 100 ms. Merely 0.004% of total stop trials had SSD greater than 600 ms, which were removed. In the first (1 -100 ms) and the last (501 - 600 ms) bin, merely 3.571 % and 28.571% of HC participants and 14.286 % and 16.327 % of TD participants respectively, contributed trials. As we had no data for a reasonable number of participants, we merged the first two and the last two bins. Thus, bins were 1-200 ms, 201-300 ms, 301-400 ms, and 401-600 ms. We labeled these bins as 1^st^, 2^nd^, 3^rd,^ and 4^th^ bins in order. We plotted the mean percentage of noncancelled responses against the midpoint of SSD, each bin for both HC and TD (Figure 2A). One-way ANOVA with SSD as a repeated factor showed a significant effect of SSD on error in inhibition in both HC [*F* (3,55) = 14.628, *p* < 0.001, 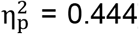] and TD group [*F* (3,98) = 47.261, *p* < 0.001, 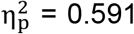]. All pairwise multiple comparison procedures (Holm-Sidak method) showed that mean error significantly increased from 1^st^ to 2^nd^ (*p* = 0.003) and from 3^rd^ to 4^th^ bin (*p* = 0.016) but a non-significant increase from 2^nd^ to 3^rd^ bin (*p* = 0.158) in HC group. Whereas the mean error increased significantly from the mean error in the preceding bin for all possible pairs (all *ps* <= 0.008) in the TD group. It shows the error monotonically increased as predicted by the race model in both groups (except error from 2^nd^ to 3^rd^ bin in HC).

**Table 1.** Behavioural performance, SSRT, and CRTT metrics in HC and TD. For both groups, average no-stop accuracy, error in inhibition, stop-signal delay, no-stop RT, and noncancelled RT are shown in the table. None of these parameters was significantly different between HC and TD. SSRT estimated by three methods was not significantly longer in TD than that of in HC group. Log-attenuation rate was significantly less in TD than that of the HC group, and the proactive delay was nearly equal. SSRT was also not significantly different between HC and TD. The standard error of the respective means (± SEM) is given in the brackets.

|  | Parameters | HC | TD | t | p |
| --- | --- | --- | --- | --- | --- |
| Behavioural performance | Go accuracy (%) | 97.09 (± 1.20) | 98.22(±0.53) | $t(75)=-0.991$ | 0.325 |
| | Stop error (%) | 42.426(±1.239) | 44.414(±0.817) | $t(75)=1.391$ | 0.084 |
| | SSD (ms) | 326(±12) | 312(±10) | $t(75)=0.81$ | 0.210 |
| | No stop RT (ms) | 556 (±13) | 540 (±9) | $t(75) = 1.04$ | 0.302 |
| | Error RT (ms) | 476 (±12) | 462 (±9) | $t(75) = 0.95$ | 0.346 |
| SSRT | Integration method (ms) | 207(±6) | 208 (±4) | $t(74)=0.241$ | 0.405 |
| | Median method (ms) | 227(±4) | 225(±4) | $t(74)=-0.36$ | 0.64 |
| | Mean method (ms) | 231(±5) | 228(±4) | $t(75)=-0.424$ | 0.664 |
| CRTT | Log-attenuation rate | 0.0097 (± 0.0003) | 0.0088 (± 0.0002) | $t(75)=-2.602$ | 0.006 |
| | Proactive delay (ms) | 389 (± 12) | 384 (± 9) | $t(75)=-0.342$ | 0.367 |

**Figure. 2.**
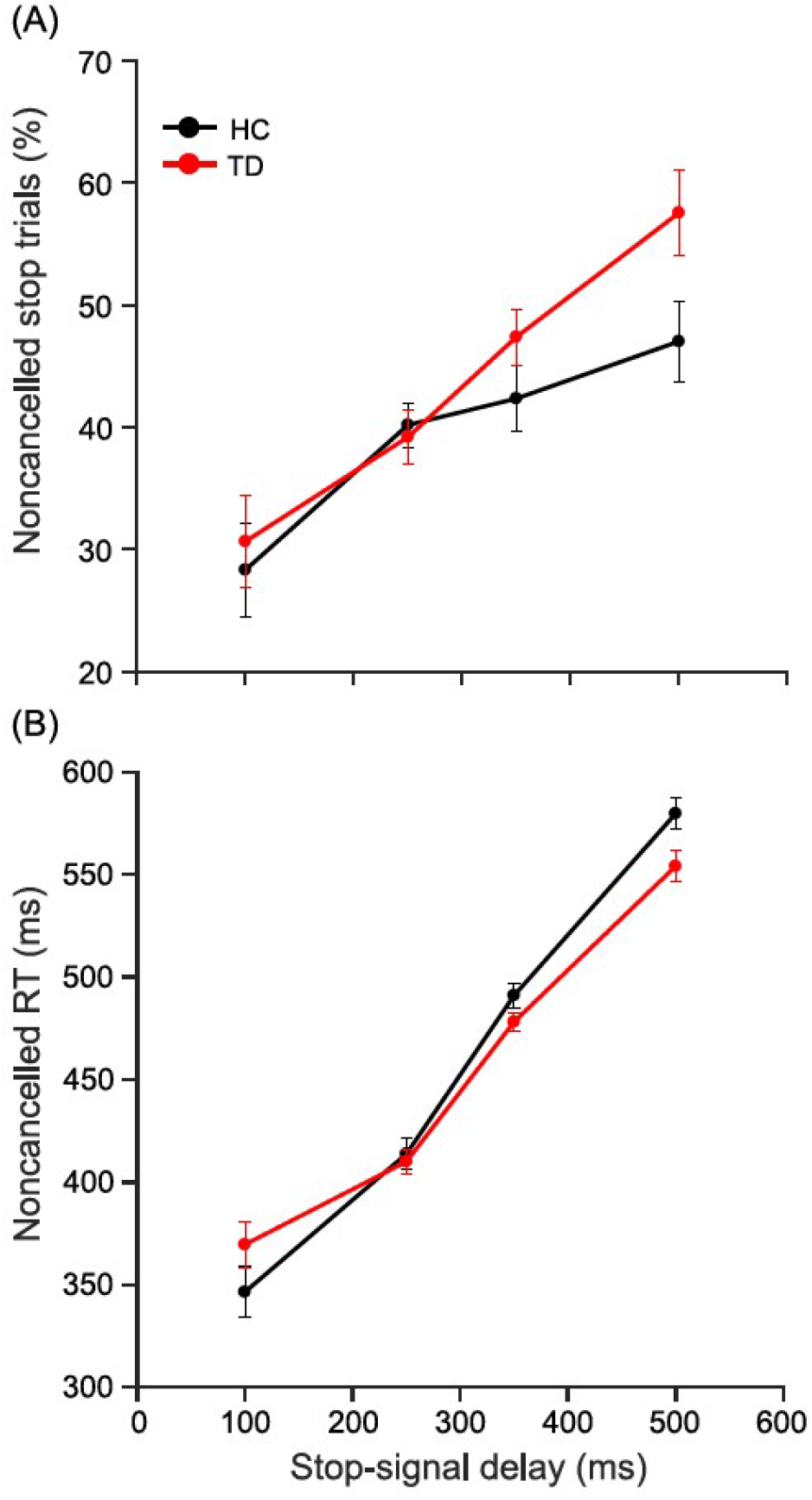
Inhibition function, noncancelled RT as a function of SSD. SSD was grouped into four (1-200 ms, 201-300 ms, 301-400 ms, and 401-600 ms) bins. The average error in inhibition and noncancelled RT in each bin for every participant was calculated. **(A)** The mean (± SEM) error in inhibition across participants for both groups (HC: black; TD: red) was plotted against the midpoint of the corresponding bin (mean: line; SEM: patch). The average percentage error in inhibition increased monotonically as a function of SSD in both groups (except a non-significant increase in error from 2^nd^ to 3^rd^ bin in HC). **(B)** The mean (± SEM) noncancelled RT across participants for both groups (HC: black; TD: red) was plotted against the midpoint of the corresponding bin (mean: line; SEM: patch). The noncancelled RT increased monotonically as a function of SSD in both groups.

### Reaction time

No-stop RT was significantly longer than non-cancelled stop RT in both HC [*t* (27) = 22.03, *p* < 0.001, *d* = 4.164, BF_+0_ >100] and TD [*t* (48) = 24.39, *p* < 0.001, *d* = 3.485, BF_+0_ >100]. Subsequently, to test whether noncancelled RT increased with the increase in SSD, we plotted mean noncancelled RT against the midpoint of each SSD bin (Figure 2B). One-way ANOVA with SSD as repeated factor showed a significant effect of SSD on noncancelled RT in both HC [*F* (3,54) = 129.717, *p* < 0.001, 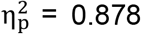] and TD group [*F* (3,95) = 95.29, *p* < 0.001, 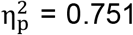]. Within both groups, mean noncancelled RT was longer than the RT in the preceding bin in all possible pairs (all *ps* <= 0.009) reflecting the replication of the predictions of the race model. In short, noncancelled RT increased monotonically as a function of SSD in both groups as predicted by the race model. These results were majorly in line with the race model’s prediction allowing estimation of SSRT.

### Stop-signal response time

We estimated SSRT by three methods to rule out any bias due to the choice of estimation method. Using the three previously described SSRT estimations (i.e., by integration, median, and mean), we found no significant difference in average SSRT between TD and HC (see Table 1). Subsequently, we checked whether and how strongly SSRT correlates with overall error in inhibition. In both HC and TD, SSRT estimated by the integration method was strongly and significantly correlated with error in inhibition, HC [*r* (52) = 0.632, *p* (one-tailed) < 0.001, *R*^*2*^ = 0.399, BF_+0_ >100], TD [*r* (92) = 0.656, *p* (one-tailed) < 0.001, *R*^*2*^ = 0.430, BF_+0_ >100]. But SSRT estimated by median method had weaker positive significant correlation with error in HC [r (50) = 0.331, p (one-tailed) = 0.049; R^2^ = 0.11, BF_+0_ = 1.673] and non-significant in TD [*r* (94) = 0.003, *p* (one-tailed) = 0.491, *R*^*2*^ < 0.001, BF_0+_ = 5.46]. SSRT by mean method was also weakly positively correlated in HC [*r* (52) = 0.43, *p* (one-tailed) = 0.013; *R*^*2*^ = 0.185, BF_+0_ = 5.096] and TD [*r* (94) = 0.248, *p* (one-tailed) = 0.044, *R*^*2*^= 0.062, BF_+0_ = 1.4]. A significant positive correlation between SSRT and error in inhibition demonstrates that the estimated SSRT was in compliance with race model. Notwithstanding that, the model turned out to be inadequate to explain the observed difference in stopping behaviour between TD and HC since there was no difference in SSRT between these groups.

### Cancellable rise-to-threshold metrics

Using CRTT metrics, we found that the log-attenuation rate was significantly less in TD than HC [t (75) = -2.602, p (one-tailed) = 0.006, d = -0.616] (Figure 3C). This difference was further supported by Bayesian t-test with ‘substantial’ evidence (BF_-0_ = 8.283). However, no significant difference was observed for the proactive delay [t (75) = -0.342, p (one-tailed) = 0.367, d = -0.081] (Figure 3D), supported by Bayesian t-test with ‘substantial’ evidence (BF_0-_= 3.123). These results suggest that TD patients’ group had a deficit in reactive control, but proactive control was intact, in comparison to the HC group.

**Figure. 3.**
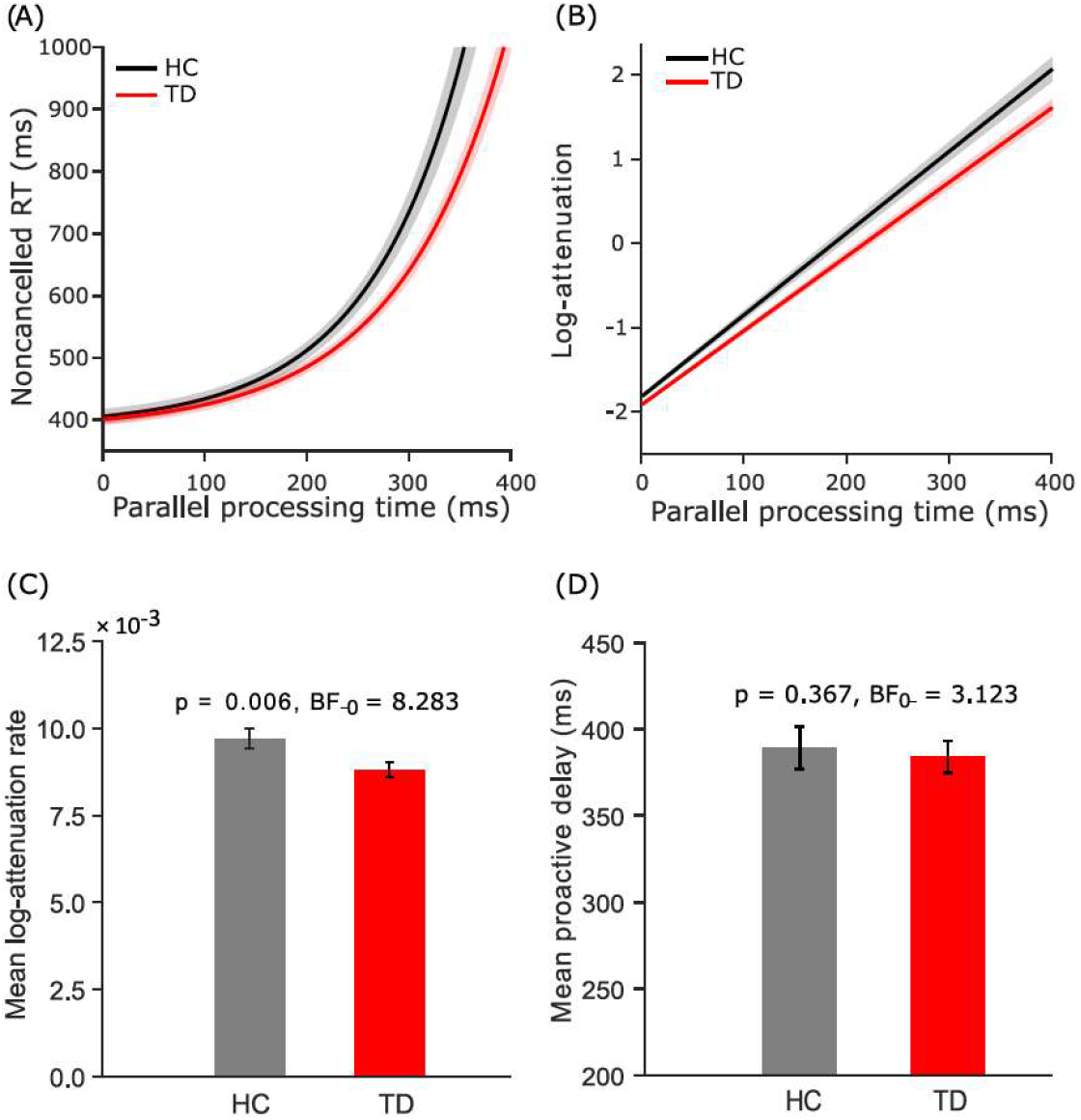
CRTT metrics between TD and HC. **(A)** The Average (± SEM) of best across participants in both groups (HC: black; TD: red) are plotted as a function of PPT (average: line; SEM: patch). It shows that there was an overall exponential increase in noncancelled RT in both HC and TD groups. The intercept at the ordinate for each fit in both groups was calculated which mathematically equals ε +c. It is referred to as CRTT metric proactive delay. Since ε was the jitter in the measurement in RT that depended on the refresh rate of the display, c was used as a proactive delay. Mean proactive delay was nearly equal in both groups. **(B)** Log of attenuation was calculated by the equation: *log*(*attenuation*) = log(*εb*) + *b* × *PPT*, at 1 ms steps from 0 to 400 ms of PPT. The average (± SEM) log-attenuation for both groups (HC: black; TD: red) is plotted as a function of PPT (average: line; SEM: patch). **(C)** The slope of the individual log-attenuation plot is referred to as the log-attenuation rate CRTT metric. The average log-attenuation in the TD group was significantly lower than that of the HC group. The mean (± SEM) log-attenuation rate (mean: bar; SEM: error bar) was plotted for both groups (HC: grey; TD: red). It shows that the TD has a significantly less log-attenuation rate than the rate in the HC. **(D)** The mean (± SEM) proactive delay (mean: bar; SEM: error bar) was plotted for both groups (HC: grey; TD: red). It shows that the proactive delay was comparable between the groups.

### Correlation of CRTT Metrics with Error in Inhibition

Theoretically, log-attenuation rate and proactive delay, each should have a negative correlation with the overall error in inhibition. We tested the correlation of log-attenuation rate and proactive delay with overall error in inhibition for both groups. Log-attenuation rate was significantly correlated with error in both HC [*r* (52) = -0.514 *p* (one-tailed) = 0.003, *R*^*2*^ = 0.264, BF_-0_ = 16.183] and TD [*r* (90) = -0.247, *p* (one-tailed) = 0.049, *R*^*2*^ = 0.058, BF_-0_ = 1.310]. However, Bayesian statistics suggests this relationship was ‘anecdotal’ (1 ≤ BF <3) for TD, but ‘strong’ (10 ≤ BF <100) for HC. Pearson’s coefficient indicates that the strength of coupling between log-attenuation rate and error in HC was almost twice the strength of coupling in TD. Proactive delay was also significantly correlated with error in both HC [*r* (52) = -0.775, *p* (one-tailed) < 0.001, *R*^*2*^ = 0.601, BF_-0_ > 100] and TD [*r* (90) = -0.735, *p* (one-tailed) <0.001, *R*^*2*^ = 0.540, BF_-0_ > 100]. Bayesian statistics suggests this relationship was ‘decisive’ (BF ≥ 100) for both groups.

### Influence of ADHD and medication on CRTT metrics

Since we found a link between deceleration rate and attention in Indrajeet and Ray (2019) and effect of medication on SSRT in Atkinson-Clement *et al*. (2020), we subsequently verified whether presence of ADHD or medication influenced CRTT metrics. We made the following observations: (1) The mean (±SEM) log-attenuation rate in TD with ADHD (0.0087 ± 0.0004) and TD without ADHD (0.0089 ± 0.0002) was not significantly different [*t* (47) = -0.364, *p* (two-tailed) = 0.717, *d* = -0.106, BF_01_ = 3.279]. (2) The mean (±SEM) proactive delay in TD with ADHD (384 ± 15 ms) and TD without ADHD (384 ± 12 ms) was not significantly different [*t* (47) = 0.003, *p* (two-tailed) =0.997, *d* < 0.001, BF_01_ = 3.461]. (3) The mean (±SEM) inhibition error in TD with ADHD (44.07 ± 1.42 %) and TD without ADHD (44.65 ± 0.99 %) was not significantly different [*t* (47) = -0.343, *p* (two-tailed) = 0.733, *d* = -0.1, BF_01_ = 3.299]. (4) The mean (±SEM) log-attenuation rate in medicated TD (0.0086 ± 0.0003) and un-medicated TD (0.0089 ± 0.0003) was not significantly different [*t* (47) = -0.828, *p* (two-tailed) = 0.412, *d* = -0.243, BF_01_ = 2.601]. (5) The mean (±SEM) proactive delay in medicated TD (366 ± 16 ms) and un-medicated TD (395 ± 11 ms) was not significantly different [*t* (47) = -1.596, *p* (two-tailed) = 0.117, *d* = -0.468, BF_01_ = 1.235]. (6) The mean (±SEM) inhibition error in medicated TD (45.89 ± 1.51 %) and un-medicated TD (43.48 ± 0.91 %) was not significantly different [*t* (47) = 1.452, *p* (two-tailed) = 0.153, *d* = 0.426, BF_01_ = 1.471]. These results indicate that the difference found in log-attenuation rate and proactive delay in HC and TD might not be due to medication or presence of ADHD. We did not consider OCD due to small sample size (N = 11) and lack of prior link between CRTT metrics and OCD.

## DISCUSSION

In this study, we applied metrics derived from both traditional race model (*1*) and relatively recent CRTT model (*27*) to assess the inhibitory control in TD patients in comparison to HC participants. Not surprisingly, like earlier studies we also observed that neither stopping behavior nor race model was able to distinguish between TD and HC. On the contrary, our new method based on CRTT model teased apart contributions from proactive and reactive control in stopping a manual movement. While the ability to procrastinate an action by TD was comparable to HC, the log-attenuation rate that estimates the strength of deceleration in GO process following the stop signal was significantly less in TD than HC. Our study also suggests that the CRTT metrics, which were originally derived to study countermanding saccades, can be generalized to variants of stop-signal task with different effectors (eye / hand) and stop-signal adjustment methods (randomization / staircasing). These results are consistent with a recent meta-analysis study indicating possible impairment in reactive but not in proactive inhibitory control in TD (*28*), and critical from the clinical point of view to identify damage in neural circuitry causing deficiency in one type of control sparing the other.

What might be a feasible physiological explanation of weak attenuation on GO process, but intact ability of strategic adjustment of the time of process initiation in the TD group? In one hand, studies have advocated that the strength of the fronto-basal ganglia connectivity largely determines the efficacy of the inhibitory control (*29, 30, 39, 31*–*38*); on the other, dysfunctional cortico-basal ganglia pathways underlie TD (*12, 40*–*43*). Interestingly, inhibition of preplanned action entails an engagement of fronto-cortical basal ganglia pathways (*33*–*39*), which is somehow compromised in TD (*12, 42, 43*). A simplified description suggests that on the direct pathway in basal ganglia, the dorsal striatum that receives projection from the frontal cortex inhibits the substantia nigra pars reticulata (SNr) causing disinhibition of the thalamus, which is attributed to an underlying neural mechanism of the GO process, whereas the subthalamic nucleus (STN) exerts an excitatory influence on SNr acting similar to pressing the brake in a car (*38, 39, 44, 45*). The dorsal striatum consists of two structures: caudate nucleus (CN) and putamen (Pu). Striatum and STN are also interconnected via globus pallidus external (GPe) on the indirect pathway (*38, 46, 47*). Imaging studies have found an abnormality in cortico-basal ganglia-thalamocortical motor loop and supplementary motor area in TD (*11, 48*). TD patients exhibited increased activation in the right supplementary motor area and the inferior parietal cortex, but decreased activation in the dorsal pre-motor cortex while performing a stop-signal task (*11, 49*). Furthermore, Ganos et al. (2014) showed that the activation of supplementary motor area (SMA) on correct stop trials were correlated with the frequency of tics. Similarly, EEG studies have also shown diminished amplitude of fronto-central P300, a marker of response inhibition, and the error positivity, while the amplitude of the error-related negativity (ERN) was elevated (*50*–*53*). Despite the differential neural activity in brain areas critical for response inhibition, none of these studies found a significant difference in SSRT or other behavioural performance between TD and HC. At the structural level, the volume of the caudate nucleus (CN), putamen (Pu), and globus pallidus (GPe) are reduced in TD adults (*43, 54, 55*). Single neuron’s activity recordings have shown a link between neuronal activity in the caudate nucleus in monkeys (*56*) and GPe neurons in rats (*38, 44*) in reactive inhibition. GPe neurons activity in rats is also linked to proactive control during stop-signal tasks (*57*). We speculate that some of the differences found at the neural level in TD might be the foundation of CRTT framework.

Our study suffers a few limitations. For example, CRTT metrics could not be computed for each emotion category and for each participant due to a smaller number of noncancelled trials in each emotion condition. It would be interesting to test whether CRTT metrics are sensitive enough to detect modulation of the inhibitory process in an emotional context and able to resolve the contradictory findings regarding the effect of emotion on inhibitory control. In addition, like many other clinical sample studies, comorbidities and medication might be issues. Although we found that CRTT metrics were not different between TD with and without ADHD, or un-medicated and medicated TD. Having said that caution should be taken interpreting no statistical difference, because effect sizes were analyzed separately ignoring the effect of other factors. Despite these limitations, our study is a methodological advancement in the assessment of deficits in inhibitory control in TD.

## MATERIALS AND METHODS

### Participants

A total of 63 TD and 34 HC were recruited for the original project. The HC group was matched with the TD group for age, gender, and acquired education, keeping the ratio between the healthy control and TD patients ∼ 1:2. The exclusion criteria for all participants were based on satisfying any of the following conditions: (1) incapable to provide consent, (2) history of substance (except nicotine) addiction or psychosis, (3) neurological condition except tics, (4) history of any psychiatric condition for HC. All participants were informed about the procedure and protocol of the study and gave their informed consent. Monetary compensation (50 Euros) was provided for participation.

### Clinical measures

All participants were assessed using the Mini-International Neuropsychiatric Interview (MINI) (*58*) for psychiatric disorders, the Minnesota Impulse Disorders Interview (MIDI) (*59*) for impulse-control related disorders, and the Barratt Impulsivity Scale (BIS)-11 (*60*) for impulsivity. For TD patients, the Yale Global Tic Severity Scale (YGTSS) (*61*) was used to assess the severity of tics. Each patient underwent a psychiatric evaluation and a medical history check-up to assess for psychiatric comorbidities.

### Stimuli and Task

As shown in Figure 1, each trial began with a white fixation cross for 500 ms at the centre of the display. It was followed by an image of a scene with emotional content for the duration of 700 ms at the centre. After a blank black display for a 1-2 s from the image, letter ‘O’ in green appeared at the centre of black display acting as a go-signal (go-trials). Participants were instructed to press the ENTER key ‘as soon as possible’ in these trials. However, in 33 % of the total trials, after a variable delay from the go-signal (stop-signal delay), letter ‘X’ painted in red appeared at the centre acting as a stop-signal (stop-trials). Participants were instructed to refrain from pressing the Enter key in these trials. For a detailed description of the task e.g., staircasing of SSD, refer to supplementary material. Details of scene image selection and categorization, and inhibitory control in induced emotional context can be found elsewhere (*12*)

### Apparatus and Procedure

E-prime software (Psychology Software Tools, Pittsburgh, PA) created and displayed all stimuli and stored the critical task parameters and responses made by participants. All stimuli were displayed on a monitor at a 60 Hz refresh rate. The ENTER key of the keyboard was assigned for response input. The pre-processing and analysis of the data was carried out by using Matlab^®^ (The Mathworks Inc., USA). Matlab^®^ Statistical Toolbox (The Mathworks Inc., USA), JASP software (JASP team, 2019, version 0.11.1) or SigmaStat (SYSTAT Inc., USA) was used for statistical analysis. Curve Fitting Toolbox™ of Matlab^®^ was used for fitting curves.

### Estimation of CRTT metrics

The CRTT model advocated that the preparatory build-up activity is decelerated after the stop-signal is detected if and only if the stop-signal is detected in a stipulated time and the magnitude of deceleration is sufficient to prevent the preparatory activity building up over time from reaching a threshold (*26*). We previously extended the CRTT model to show an exponential increase in reaction time (RT) in unsuccessful stop trials, as parallel processing time (PPT) increased (*22*). PPT is defined as the maximum time duration, for which the stop-signal and the go signal can be processed in parallel before the elicitation of a response. Empirically, it was calculated by subtracting SSD from RT in error stop trials (i.e., noncancelled RT – SSD). PPT was grouped in bins and mean PPT and corresponding mean noncancelled RT was calculated for each bin. Noncancelled RT was plotted against PPT, and the plot was fitted with an exponential function (noncancelled RT =ε e ^b(PPT)^ +c). The rate of log of the slope of the best exponential fit is referred to as log-attenuation rate, and the intercept of the best exponential fit is referred to as proactive delay. For a detailed derivation of CRTT metrics, refer to supplementary material or our previous work (*27*).

### Statistical analyses

The criterion of significance α was fixed at 0.05 level, except for the tests of normality and equivalence of variance where a lenient criterion was used (α = 0.01). Linear Pearson product–moment correlation method was performed to calculate correlations. (see **supplementary materials** for more details).

## Author contributions

PP, YW conceived the study, and designed the experiment; CA conducted experiment and collected data; II analyzed data; SR designed the model; all authors wrote the manuscript.

## Data and materials availability

Data may be shared in its raw form for review purpose or other reasonable scholarly use.

## Supplementary Materials for

### Supplementary Text

#### Emotional Stop Signal Task (ESST)

After a variable interval between 1-2 s from the image, the letter O in upper case and painted in green color appeared at the center of the black screen, which acted as a *go-signal* instructing the participant to press the ENTER key of the keyboard *“as soon as possible”. Maximum* allowed period to press the key was 1000 ms, otherwise, the trial was aborted. The duration between the onset of the go-signal and the keypress is referred to as go (or no-stop) reaction time. In 33 %of all trials, called *stop trials*, after a variable delay from the onset of the circle, the letter X in upper case and painted in red color replaced the letter O at the center. The appearance of the letter X acted as the *stop-signal* instructing participants to refrain from pressing the key. The delay between onset of the go- and stop-signal is called stop-signal delay (SSD), and it was adjusted by the tracking procedure. In this procedure, initially, the SSD was fixed at 250 ms in each emotion category. Subsequently, SSD was increased by 25 ms, if the previous stop trial was correct (i.e., the response was cancelled); otherwise, SSD was decreased by 25 ms. In the rest of 66 %of trials, called *go-trials* or *no-stop trials*, the stop-signal did not appear, and participants were required to press the ENTER key. No-stop and stop trials were randomly interleaved. No feedback about the outcome (correct or incorrect) of a trial was provided. However, if the go response was too slow (> 1000 ms), a message requesting to generate a faster response in the next trials was displayed.

#### CRTT metrics

We defined PPT as the maximum time duration, for which the stop-signal and the go signal were processed in parallel before the elicitation of a response. Empirically, it was calculated by subtracting SSD from RT in error stop trials (i.e., noncancelled RT – SSD). When the stop-signal appeared after the response was generated (PPT<0), it could not influence the ongoing motor plan. Therefore, only a subset of trials that yielded PPT ≥0 was used for the estimation of CRTT metrics. We estimated these metrics by fitting noncancelled RT and PPT data with an exponential function (noncancelled RT = ε e^b(PPT)^ +c). We previously derived CRTT metrics to assess the efficacy of inhibitory control in HC (*27*). In brief, the rate of change of non-canceled RT concerning PPT gives the slope of the function RT =ε e ^b(PPT)^ +c, which estimates attenuation exerted on GO process building up over time to reach a decision threshold as a function of PPT [*attenuation* =εb e ^b(PPT)^]. After taking the natural log of both sides we get, ln(*attenuation*) =ln (*εb*) + *b* × *PPT*. Alternatively,ln(*attenuation*) = *b* × *PPT* + *b*_O_, where b_0_ equals ln (*εb*). Given that *b* is obtained from the fitting algorithm and fixed for each participant, and *ε* is a fixed nominal error in the estimation of RT, the rate of change in attenuation in log-scale. Concerning PPT equals to We refer to this fitting coefficient as ‘log-attenuation rate’, which is one of the CRTT metrics. Whereas the other CRTT metric, which we refer to as ‘proactively delay’, estimates participant’s ability to procrastinate response elicitation in anticipation of the stop-signal. Proactive delay is calculated by inserting PPT= 0 in RT =ε e ^b(PPT)^ +c. Since ε is constant and nominal in comparison to noncancelled RT, (ε +c) is approximated to the fitting coefficient *c* to estimate proactive delay.

#### Statistics

We first compared behavioral performance (e.g., average error and RT in go and stop trials) in stop-signal tasks and between HC and TD using t-tests. We used one-way repeated measures ANOVA to see whether stop error and RT in error stop trials increased with SSD. Holm-Sidak method was used for all pairwise comparisons wherever required. Subsequently, we compared SSRT between TD and HC estimated by three methods by t-tests and checked the correlation of SSRT with average error in inhibition. Next, we fitted the CRTT model to the plots of noncancelled RT against PPT for both groups. Bayesian versions of tests with recommended (e.g., default prior) settings in JASP were performed and Bayes factor (BF) is reported that was specifically aimed for null results’ interpretation. With the JASP default value of the r-factor (0.707), we fixed criteria of BF values for classifying strength as anecdotal (1 ≤ BF <3), substantial: 3 ≤ BF< 10, strong: 10 ≤ BF <100, and decisive: BF ≥ 100.

#### CRTT model fitting and outlier criteria

Since we did not have a reasonable amount of data for each emotion category for CRTT model fitting, we could not take emotion into account. We grouped noncancelled stop trials of each participant from 0 ms to 400 ms in bins of size 80 ms. The mean PPT and mean noncancelled RT were calculated across trials in each bin. The mean noncancelled RT was plotted against the mean PPT for each participant and fitted with the exponential function (noncancelled RT = ε e ^b(PPT)^ +c). We fixed ε at 17 ms which was almost equal to one refresh duration of the display monitor to account for random jitter in the measurements of RT and SSD. In the HC, one participant had data only in two bins of PPT, which was less than the minimum data points required to fit it to the exponential function, so the participants’ data were removed. The goodness-of-fit for two participants, one in each group (HC: *R*^*2*^ = 0.26, TD: *R*^*2*^ = 0.001), were less than the minimum fixed criterion (*R*^*2*^ < 0.5). Besides, we used the box-and-whisker plot method or Tukey method (*62*) to detect univariate outliers, if any, in b or c in both groups. Three participants from the TD group were marked outlier by the box-plot method (b values: 0.0130, 0.0129, and 0.0130, number of IQR from median: 1.9979, 1.93, and 1.978). No univariate outliers were detected in c in the TD group. No outliers were detected in b or c in the HC group. Thus, two participants’ data from HC and four from the TD group were removed as outliers from all subsequent analyses. *The final sample contains 28 HC and 49 TD*. Besides, in the HC group, one participant had an SSRT of 19 ms, which is theoretically not possible, so it was removed from wherever SSRT of the HC was used. In correlation analyses, scatter plots with fitted regression lines with 95 %confidence bound boundaries were used. Any data points falling out of the prediction bound were removed as outliers from the correlation analysis.

